# The relationship of survivorship and body mass modeled by metabolic and vitality theories

**DOI:** 10.1101/129809

**Authors:** James J. Anderson

## Abstract

The relationship between body mass and survivorship is explained by a model that merges metabolic theory relating metabolism to body mass, and vitality theory relating survival to vitality loss and extrinsic mortality. The resulting metabolic-vitality framework hypothesizes mortality results from replicative senescence of the hematopoietic system and predator-prey interactions. Fitting the metabolic-vitality model to body mass and maximum lifespan data of 494 nonvolant mammals yields allometric relationships of body mass to the vitality parameters, from which full survivorship profiles can be predicted from body mass. Comparisons of the mass-derived vitality parameters to those estimated directly from survival data identifies how intrinsic and extrinsic mortality processes of specific populations deviate from the aggregate. Highlighted findings include a mathematical explanation for the shift from Type I to Type II survivorship curves with decreasing body mass, a quantification of the impact of hunting on wild populations and a quantification of the reduce rate of primate aging relative to the aggregate of mammal populations. Finally, the framework allows explorations of the combined effects of animal aging and predation on survival patterns.

## Introduction

The relationships of body size to animal physiology, metabolism, life history and ecology (Blueweiss et al. 1978; Calder 1984; Lindstedt and Calder 1981) convey important information on the processes shaping the evolution and functioning of the biosphere. Most important of these is the quarter-power scaling of metabolism *B* to body size in mass *M*, *B* = *aM*^3/4^, (Kleiber 1947) known as the WBE Theory or simply metabolic theory (Banavar et al. 2010; Brown et al. 2004; West et al. 1997). In the theory the ¾ exponent emerges by the limitation on metabolism by the rate materials move through the branching networks of an animal’s circulation system. The theory has also been extended to life history (Brown et al. 2004) and a metabolic theory of ecology (Humphries and McCann 2014). The relationship of body size to metabolism also plays a central role in the study of evolution and life histories of extinct species. In particular, Cope’s rule (Cope 1904), in which over evolutionary time the body size of clades increased by orders of magnitude (Alroy 1998; Benson et al. 2014; Evans et al. 2012; Smith et al. 2010), is thought to be driven by interactions of ecological process and abiotic forcing (Raia et al. 2012; Saarinen et al. 2014; Segura et al. 2016). However, the body size increase also may be controlled by intrinsic biological factors (Sookias et al. 2012) that are limited by the allometric relationships of mass to metabolism and generation time, which in turn set the rate of gene mutation (Okie et al. 2013).

Thus, critical to extending the relationships of body size to life history and evolution is the relationship of body size to animal lifespan and survivorship. While an approximate quarter-power relationship of body size to maximum lifespan is well established (Blueweiss et al. 1978; de Magalhães and Costa 2009; de Magalhães et al. 2007; Lindstedt and Calder 1981), a mechanistic model underlying the relationship have not been elaborated and the shape of survivorship curves with body size has not been addressed. Part of the difficulty involves the fact that mortality results from both intrinsic aging processes and environmental challenges. Intrinsic factors that involve cellular level aging should be implicitly connected to body size through metabolism. Extrinsic factors may also involve body size through the ability of larger animals to avoid predators. Additionally, mode-of-life effects the rate of mortality, with ability to fly being of particular importance in explaining the longer lifespan of birds relative to mammals (Healy et al. 2014).

The goal of this paper is to develop a mathematical model linking body size to longevity through cellular level processes. The framework combines metabolic theory (Brown et al. 2004; West et al. 1997), processes of cell aging (López-Otín et al. 2013) and vitality theory (Anderson 2000; Anderson et al. 2017; Li and Anderson 2013).

## Model

### Metabolic theory

The model begins with metabolic theory that characterizes Kleiber’s law in terms of the constraints on the distribution rate of resources to cells. The law expresses a power-law relationship between metabolism and body size as

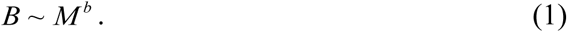

where *b* takes on values approximately ∼3/4^1^ The relationship emerges from the assumption of the minimization of energy dissipation in a fractal network of branching tubes, i.e. the circulatory system that distributes blood from the heart to cells through the vascular network (Brown et al. 2004; West et al. 1997). A further extension of the theory invokes a cascade of power-relationships between body size and metabolism that result in the observed power-law. The processes, which together control oxygen supply to mitochondria for the synthesis of ATP, thus work on scales of molecules to organs (Darveau et al. 2002).

These allometric models predict that the amount of blood servicing a metabolic site increases with body size. To set the foundation of the model first consider the unique allometric properties of the physiological systems involved with oxygen transport. Ninety percent of the oxygen transported is used in muscle work (Darveau et al. 2002). The systems directly involved, i.e. muscle, blood, spleen, and skeleton, scale with body size at powers greater than one while the system not directly involved, the digestive, renal, integumentary, endocrine and nervous systems, scale with powers less than one (Calder 1984). As a consequence, the rate at which a blood cell supplies oxygen to a metabolic site declines with body size as ∼ *M*^−1/4^ while, the cell’s cycle time (Banavar et al. 2010) and lifespan (Gillooly et al. 2012) both increase with body size as *M*^1/4^. Taken together, these allometric relationships indicate that the demand on the individual cells of the circulatory system decline with increasing body size with the –¼ power. Specifically this allometric scaling applies to the erythrocytes (red blood cells) (Wiegel and Perelson 2004) and the leukocytes (white blood cells) that protect somatic cells from pathogens and damage. In particular, leukocytes play a key role in regulating aging^2^.

### Hematopoietic hypothesis of longevity

The central hypothesis of the model involves the relationship of leukocyte dynamics with body size and focuses on the hematopoietic system in which hematopoietic stem cells (HSC) in the bone marrow are the source of leukocytes, which are some of the most rapidly replaced cells of the body (order of days). Thus, relationships between body size, the immune system and hematopoietic stem cells (HSC) are potentially central to the body size-longevity pattern. Importantly for these relationships, the HSC population is independent of body size while the rate of production of blood cells and longevity scale in inverse manners with body size such that the number of replications of HSC is constant over lifespan and independent of body sizes (Dingli and Pacheco 2006).

Thus lifespan may have a cellular level basis through, what has been termed, the replicative senescence” hypotheses (de Magalhães and Faragher 2008). This hypothesis draws on aging research showing that interventions that induce premature aging and shortened longevity involve increased cell apoptosis and accelerated senescence, while interventions that increased longevity reduced apoptosis and delay senescence. Replicative senescence focuses on aging variations within a species, but to extend the hypothesis across species, the cell level functions need to be linked to body size. Metabolic theory provides this link.

Clearly, aging and lifespan involve more than a single physiological pathway (Kirkwood and Austad 2000; Warren and Rossi 2009) and a hierarchy of functions from the molecular processes of a cell to intercellular communication are associated with aging (López-Otín et al. 2013). Importantly, evidence indicates these functions are regulated by body size. Specifically, the metabolic rate of mammalian cells increase in isolation, becoming essentially indistinguishable from the metabolism rate of a unicelled organism (West et al. 2002). Additionally, manipulations that alter the rate of apoptosis and senescence, measures of aging, correspond with changes in lifespan and suggest that an organisms lifespan is linked to the number of times a cell divides (de Magalhães and Faragher 2008). Thus, the fixed number of HSC in mammals, their function as progenitors of leukocytes and the correlation of cell division with body size all point to the possibility that the longevity-body size relationship has its basis in the hematopoietic system. A brief discussion of hematopoietic links to lifespan follows.

#### Hematopoiesis and aging

Adult blood cells are constantly regenerated by self-renewal and hierarchal differentiation of HSC through series of progenitor stages that results in the production of erythrocyte (red) cells and a wide variety of leukocytes (white) cells (Jagannathan-Bogdan and Zon 2013; Rieger and Schroeder 2012). In humans, HSC resides in an active pool of ∼ 400 cells and a quiescent reserve of ∼ 10^4^ cells (Dingli and Pacheco 2010). The rate of replication of HSC decreases with increasing body size as *r*_*HSC*_ ∼ *M*^−1/4^ (Dingli and Pacheco 2006; Dingli et al. 2008) such that on the average a HSC divides every few weeks in a mouse and about every year in a human (Gordon et al. 2002). While the total HSC population in adult mammals is independent of body size (Abkowitz et al. 2002; Dingli et al. 2008), through self-renewal the HSC pool appears sufficient to maintain the blood supply through life which is on the order of 10^11^ blood cells per day for humans (Gordon et al. 2002). Noting that lifespan scales as approximately *M*^1/4^ and using the replication rate *r*_*HSC*_, then the total number of replications is *T* ∼ *M*^1/4^ *M*^−1/4^ ∼ *M*^0^. Thus, the total number of HSC replications over a lifespan should be independent of body size. Also of importance, *T* is on the order of the Hayflick limit^3^ (Dingli et al. 2008). However, while *T* is essentially the same across the 6 orders of body size magnitude, e.g. African pygmy mouse (10 g) to elephant (5000 kg), there is no clear evidence that HSC replicative senescence limits the production of blood or directly contributes to blood aging. However, through their slow replication HSC may accrue DNA damage, erosion of telomere end caps and accumulate epigenetic instability (Warren and Rossi 2009). Through the multiple cell transformations and replications between creating of a HSC progenitor cell to the terminal blood lineages, damage accumulates leading to reduction in the capacity of the hematopoietic system. Importantly, the damage can be inherited by different progeny cells as well as propagated through self-renewal of the stem cells (Beerman 2017). The combination of these factors implies a gradual aging of the blood system and not a sharp onset of replicative senescence. However, the finding that across the body-mass spectrum all mammals have equivalent HSC populations and the correspondence of maximum number of replications of individual stem cells to the Hayflick limit are compelling evidence for a central molecular basis of longevity linked through the body size regulation of cell metabolism.

Furthermore, the degradation of the hematopoietic system can propagate into immunosenescence and the deregulation of somatic tissue (Henry et al. 2011; Kovtonyuk et al. 2016; Park 2017), which can lead to proximal causes of mortality such as cancer (Rozhok and DeGregori 2016; Rozhok et al. 2014) and chronic stress that can feed back to increase leukocyte production resulting in atherosclerosis, inflammation and cardiovascular disease (Elias et al. 2017; Hanna and Hedrick 2014; Heidt et al. 2014). While the model focuses on the replicative limit of HSC and accumulative damage of the hematopoietic pathways, the interaction is not unidirectional. The sympathetic nervous system regulates the HSC self-renewal rate, pool size and the rate blood cells egress to the circulation system (Bellinger and Lorton 2014). Thus, the replication rate of HSC is determined from the body status as tracked by the sympathetic nervous system, and is highly tuned to the animal and its environment.

Other studies support the centrality of body size in regulating the efficiency of the hematopoietic system and immune system efficiency. At the cellular level, lymphocyte numbers scale with body size such that larger animals have more immune system protection than smaller animals (Perelson and Wiegel 2009; Wiegel and Perelson 2004). Additionally, a study of blood of 85 birds species revealed large-brained birds exhibit lower oxidative damage to lipids and had higher antioxidant capacity leading to a hypothesis of “oxidative avoidance” with large body size (Vágási et al. 2016). Notably, the hematopoietic system link to longevity does not directly involve cellular oxidative stress, but it does provide one possible explanation for how cellular level processes affect life history properties, a topic of significant interest in longevity science (Speakman et al. 2015).

### Vitality theory

To link metabolic theory, HSC dynamics and longevity first define an age-dependent survival capacity*V*_*x*_ at age *x*, as an index of the capacity of the hematopoietic system to maintain the immune system. The initial survival capacity*V*_0_ is an index of the number of HSC and is independent of body size. Therefore, the initial survival capacity is a constant across all mammal body sizes. Define a normalized survival capacity, designated vitality, as *v*_*x*_ = *V*_*x*_/*V*_0_ where *v*_0_ = 1 is the initial vitality and like the initial survival capacity, is independent of body size. Vitality declines with age from the gradual accumulation of damage and mutations in the hematopoietic system and the rate of decline is characterized by a Wiener process as

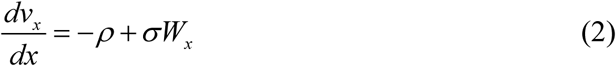

where *W*_*x*_ is a noise process with zero mean value and unit variance, *ρ* is the mean rate of loss of vitality and *σ* scales the level of stochastic variability in the rate (Anderson 2000; Anderson et al. 2017; Li and Anderson 2013). Equation (2) characterizes an ensemble of random vitality trajectories that begin at *v*_0_ = 1 and drift towards a zero boundary where they are absorbed (Fig. 1). Each path represents the probabilistic realization of an individual’s immune system function from its initial state to its end state.

In the vitality framework, the mortality results from the loss of vitality either from the stochastic degradation of vitality according to eq. (2) or from the loss of vitality from an environmental challenge to the remaining vitality (Fig. 1) (Li and Anderson 2013).

**Fig. 1.**
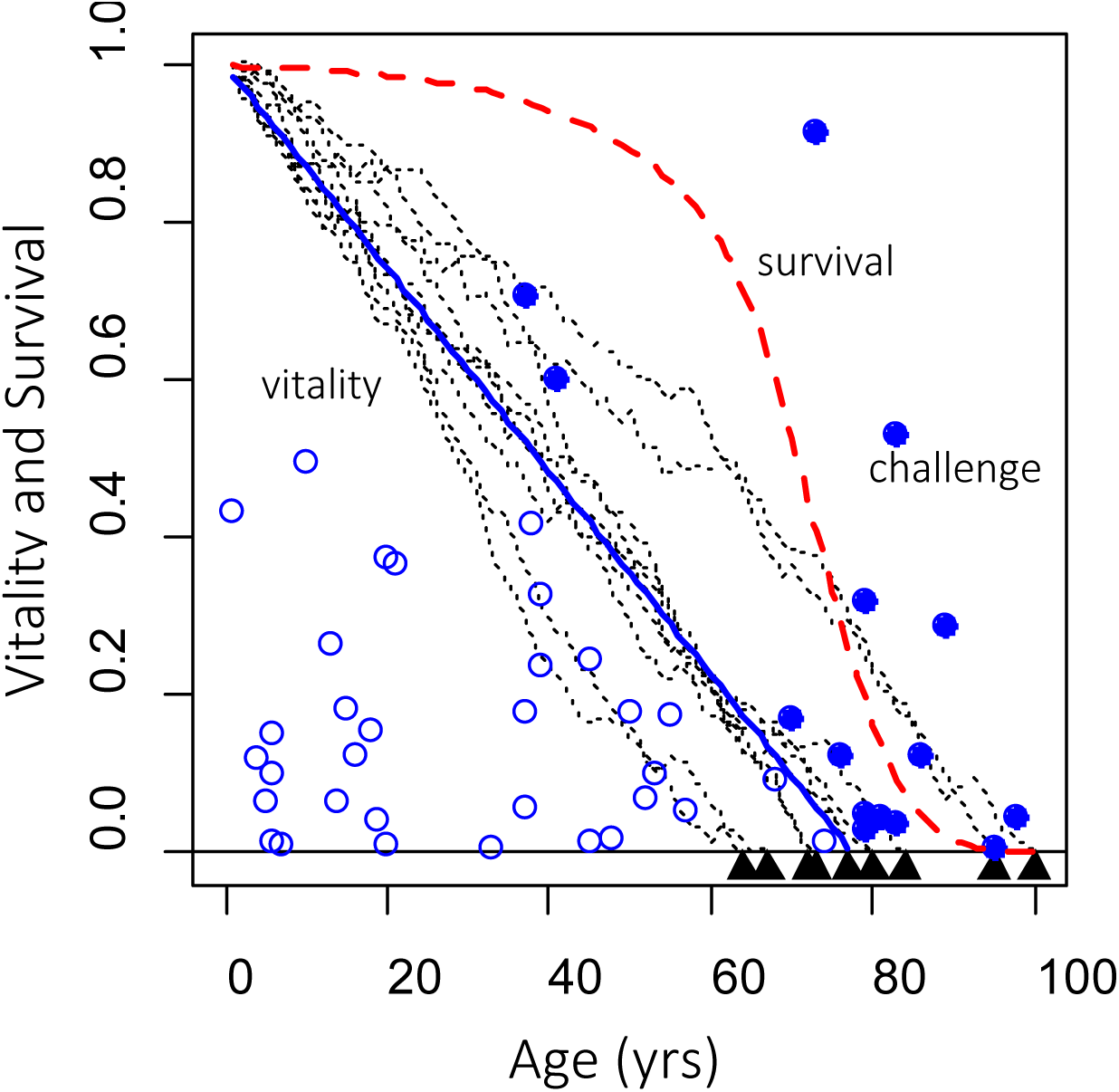
Illustration of extrinsic and intrinsic mortality processes defined by eqs (5) and (3). Extrinsic mortality results when a challenge event magnitude (circle) exceeds the age-specific population-level adult vitality (blue line). Filled blue circles indicates fatal event, open circles nonfatal events. Adult intrinsic mortality results when stochastic vitality paths reach the zero boundary. The figure was generated using *ρ* = 0.0129, *σ* = 0.0126, *λ* = 0.045, and *β* = 0.20.

#### Intrinsic survival

The rate of mortality is calculated from the probability distribution of arrival time of random vitality paths defined by eq. (2) to the zero boundary. Ignoring the preferential removal of lower vitality paths by extrinsic mortality, survival with intrinsic processes only is

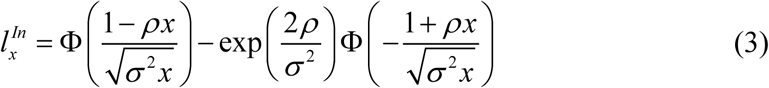

With 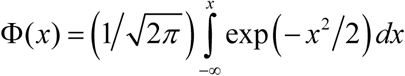 the cumulative standard normal distribution (Li and Anderson 2013).

#### Extrinsic survival

Extrinsic mortality results when an environmental challenge such as disease, predation or physical injury exceeds the age-specific survival capacity of the organism. With stochastically stable challenges, the extrinsic mortality rate increases with age as vitality *v*_*x*_ declines. To represent this process assume challenges have a Poisson frequency distribution with mean *λ* and an exponential magnitude distribution *φ*(*z*) = 1– e^− *z* / *β*^ with mean *β*. Then the conditional extrinsic mortality rate for a given vitality *v*_*x*_ is

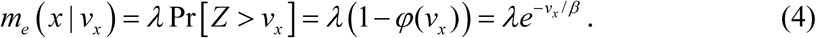

Deriving the extrinsic mortality rate requires integrating eq. (4) over the probability distribution of *v*_*x*_. Because challenges preferentially remove low-vitality individuals, the probability distribution of *v*_*x*_ trajectories defined by eq. (2) is modified and so does not have a closed form solution. However, the stochastic distribution of vitality can be approximated by the linear form *v*_*x*_ = 1– *ρx*. Then challenge events are fatal if their magnitude exceed the linear approximation of vitality (Fig. 1). The resulting extrinsic mortality rate is *u*_*e*_ = *λe*^−(1–*ρx*) *β*^ (Li and Anderson 2013). The survivorship curve from extrinsic mortality processes in the absence of intrinsic mortality is then

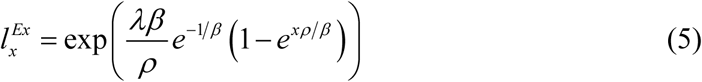

where the challenge magnitude has the range 0 ≤ *β* <∞. When *β* = 0 extrinsic mortality is zero and 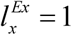. When any challenge exceeds the available vitality, i.e. *β* ≫ 1 then the extrinsic survivorship profile reduces to an exponential form 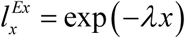. As an aside, a juvenile extrinsic mortality rate, involving challenges to a developing juvenile immune system can be expressed in a similar manner (Anderson et al. 2017). This paper does not consider juvenile extrinsic mortality.

### Total survival, *ELS, MLS*

The total survival as a function of age and expected lifespan (*ELS*) and maximum lifespan (*MLS*) are now defined

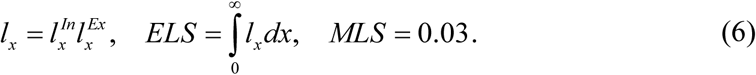

The maximum lifespan is set at 3% survival. This is an arbitrary value since the maximum lifespan is operationally determined by the mean of the oldest *N* individual independent of the initial population size. Typically *N* is above 100 individuals (de Magalhães and Costa 2009).

### Relating model coefficients to body size

To model the effect of body size on longevity the parameters *ρ*, *σ*, *β* and *λ* are expressed in terms of adult body size based on the reasoning below.

#### Intrinsic factors

Equation (2) expresses the rate of decline of vitality *ρ*. Based on the relationship of hematopoiesis to aging and metabolism discussed above, the rate of decline relates to body size as *ρ* ∼ *M* ^− *k*^ where the metabolic exponent is a free parameter, *k* ∼ 1/4. The stochastic variation in the rate is assumed to follow the same relationship, giving*σ* ∼ *M* ^−*k*^.

#### Extrinsic factors

Extrinsic mortality results from acute challenges to the remaining vitality and, in general, partitions into challenges from disease, physical injury and predation. In all cases, the magnitude and frequency of the challenges involve the state of the environment and mortality results when the challenge magnitude exceeds the vitality level, which depends on body size and age. Thus, the extrinsic parameters describe a complex of processes that may relate to body size.

To establish these relationships, consider first the frequency of challenges from pathogens and predators. Assume the frequency of encountering a deadly pathogen is independent of body mass and therefore is disregarded in the model. However, mass-independent challenges could readily be included as additional parameter. An equation for predator encounters is based on macroecological theory (Hatton et al. 2015). Assume the encounter frequency is related to the ratio of population densities (number/area) as *λ* ∼ *N*_*pred*_*/N*_*prey*_ where prey designates the species of interest. Predator-prey studies indicate that the predation rate declines as a power law with increasing prey population biomass. Across a range of large mammal ecosystems, predator and prey biomass densities (total biomass/area) exhibited the relationship 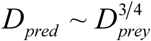 where *D* _*i*_ = *M*_*i*_ *N* _*i*_.

Furthermore, the size structure of predator and prey communities was found nearly constant across biomass gradients. That is, mean body mass of predators *M* _*pred*_ and prey *M* _*prey*_ were found to be invariant across range of prey densities *D*_*prey*_. Thus, to a first order the predator and prey body masses have a proportional relationship *M* _*pred*_ ∼ *M* _*prey*_. It follows the predation rate on the prey in a trophic structure can be expressed 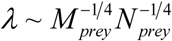. This relationship suggests predation rate decreases as a –¼ power as both prey body mass and population density increase. However, the population density is likely determined by ecological factors not clearly linked to prey or predator body masses and therefore for this model *N*_*prey*_ is assumed constant.

Addressing the challenge magnitude, the model assumes magnitudes are random and follow an exponential distribution in which low magnitude challenges occur more often than high magnitude ones. Furthermore, because challenges originate from the environment, assume their magnitude and animal body size are independent. Although this is not necessarily the case, in predation the predator-prey pairing by definition means the predator can capture and kill the prey. Thus, to a first order a predator challenge exceeds the available vitality of the prey and therefore *β* ≫ 1. With this assumption, the effect of challenges on extrinsic survival is characterized by their frequency *λ* only. However, when considering the effects of challenge magnitude on survivorship patterns across time, temporal changes in challenge magnitude can be important (Anderson et al. 2017).

### Final model relating survival to body size

The final equation, characterizing the effects of intrinsic and extrinsic mortality processes on survival as a function of animal body size is

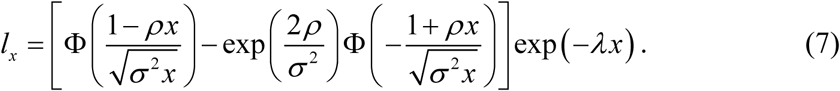

The set of parameters *Q* = [*ρ*,*σ*, *λ*] that can be estimated in two ways. Designate *Q*_*S*_ as parameters estimated by fitting eq (7) directly to survival data. Designate *Q*_*M*_ as parameters estimated by fitting body mass to maximum lifespan data using the equations

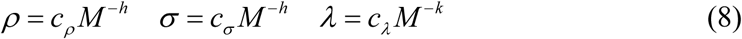

In this second case the estimation routine fits the parameter set *P* = [*c*_*ρ*_, *c*_*σ*_, *c*_*λ*_, *h*, *k*].

### Model Fitting

#### *Fitting* log *MLS vs.* log *M*

The model predicts a relationship between body size and maximum lifespan, in terms of eq. (7) with coefficients defined by eq. (8). These coefficients were estimated by fitting the model to log *MLS* to log *M* data using the R^(c)^ language nls2 nonlinear fitting package that can explore a user-defined parameter space with a random-search algorithm. The parameter space was searched with 10,000 parameter sets *P*_*i*_ = [*c*_*ρ*_, *c*_*σ*_, *c*_*λ*_, *h*, *k*]. For each set the method calculated log *MLE* using the bisection method in which beginning with initial boundaries *x*_*lower*,*i*_ < *MLS* and *x*_*upper*,*i*_ > *MLS* the algorithm calculated the midpoint *x*_*middle*,*i*_ and iteratively adjusted boundaries replacing the appropriate boundary with the previous middle point, e.g. *x*_*middle*,*i*_ < *x*_*middle*,*i*+1_ < *x*_*upper*,*i*_, until 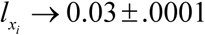. Residual sum of squares *RSS*_*i*_ = ∑ _*j*_ (log *MLS* _*j*_ – log *MLS* _*j*_ (*P*_*i*_)) were calculated for each *P*_*i*_ with the optimal *P*_*opt*_ having the minimum RSS.

The estimation procedure does not yield confidence limits. Therefore, to explore the effect of variability in parameters the search was repeated 20 times for 500 random parameter values confined to the parameter range 0.9*P*_*opt*_ to1.1*P*_*opt*_ for each repeat. The standard deviations of the parameters determined from the 20 sets of *P*_*opt*_ were taken as statistical measures of the fitting routine confidence intervals.

#### Fitting Survival data

Parameters *Q*_*S*_ were estimated from survivorship curves with a maximum-likelihood routine that fits eq. (7) to survival data (Salinger et al. 2003). The algorithm “vitality.k” is available in R code at www.CRAN.R-project.org/package=vitality. The relationships between the *Q*_*S*_ parameters and a survivorship curve shape is detailed in (Li and Anderson 2009). In essence, increasing *ρ* decreases *ELS* while increasing *σ* decreases the slope of the survival curve about the *ELS*. Increasing *λ* increases the overall curve slope with the greatest effect in early life prior to the onset of intrinsic mortality.

### Data

#### Maximum lifespan data

To estimate the *P*_*opt*_ parameters the model was fit to maximum lifespan and body weight data drawn from the date set in Appendix 2 of (Healy et al. 2014). The data which originated from the AnAge database (de Magalhães and Costa 2009), consisted of 494 nonvolant terrestrial mammal species, each constructed with over 100 maximum lifespan records.

#### Survivorship data

To characterize ability of eq. (7) to fit survivorship curves survival data was extracted from 6 published studies representing captive and wild species, different life history strategies and body sizes. Survival data for field vole (*Microtus agrestis*) (Selman et al. 2013), pigtail macaque (*Macaca nemestrina*) (Ha et al. 2000) and Asian elephant (*Elephas maximus*) (Sukumar et al. 1997) were collected from captive populations. Survival data for female mountain gorilla (*Gorilla beringei beringei*) (Robbins et al. 2011),Yellowstone grizzly bear (*Ursus arctos horribilis*) (Knight and Eberhardt 1985) and Namibian cheetah (*Acinonyx jubatus jubatus*) (Marker et al. 2003) were collected from wild habitats.

### Results

#### Fit of log *M* to log *MLS*

Table 1 shows the optimal parameter set *P*_*opt*_ and estimates of the variability and Fig 2 shows the resulting fit of the model to log body size to log maximum lifespan. The predicted relationship has a slight curvature compared to a linear regression. However, the residual-sum of squares of the model (RSS = 69.70) and linear fit (RSS = 69.65) are nearly identical. Note, the linear regression of log *MSL* vs. log *M* yields a slope of 0.19 and the best-fit model parameters, *P*_*opt*_ yields the same slope with the allometric parameter *h* = 0.24.

**Table 1.** Parameters generated by fitting the model to log *M* vs log *MLS* data. *P*_*opt*_ coefficients for terrestrial nonvolant mammals. *P*_20_ Indicates the mean of 20 runs with 500 searches of the parameter space ±10%*P*_*opt*_ for each run. SD_*P*200_ depicts the standard deviations of the runs.

| | $c\rho$ | $c\sigma$ | $c\lambda$ | $h$ | $k$ |
| --- | --- | --- | --- | --- | --- |
| $P_{opt}$ | 0.230 | 0.358 | 0.044 | 0.241 | 0.300 |
| $P_{20}$ | 0.230 | 0.366 | 0.046 | 0.241 | 0.307 |
| $SD_{P_{20}}$ | 0.005 | 0.015 | 0.002 | 0.003 | 0.013 |

**Fig. 2.**
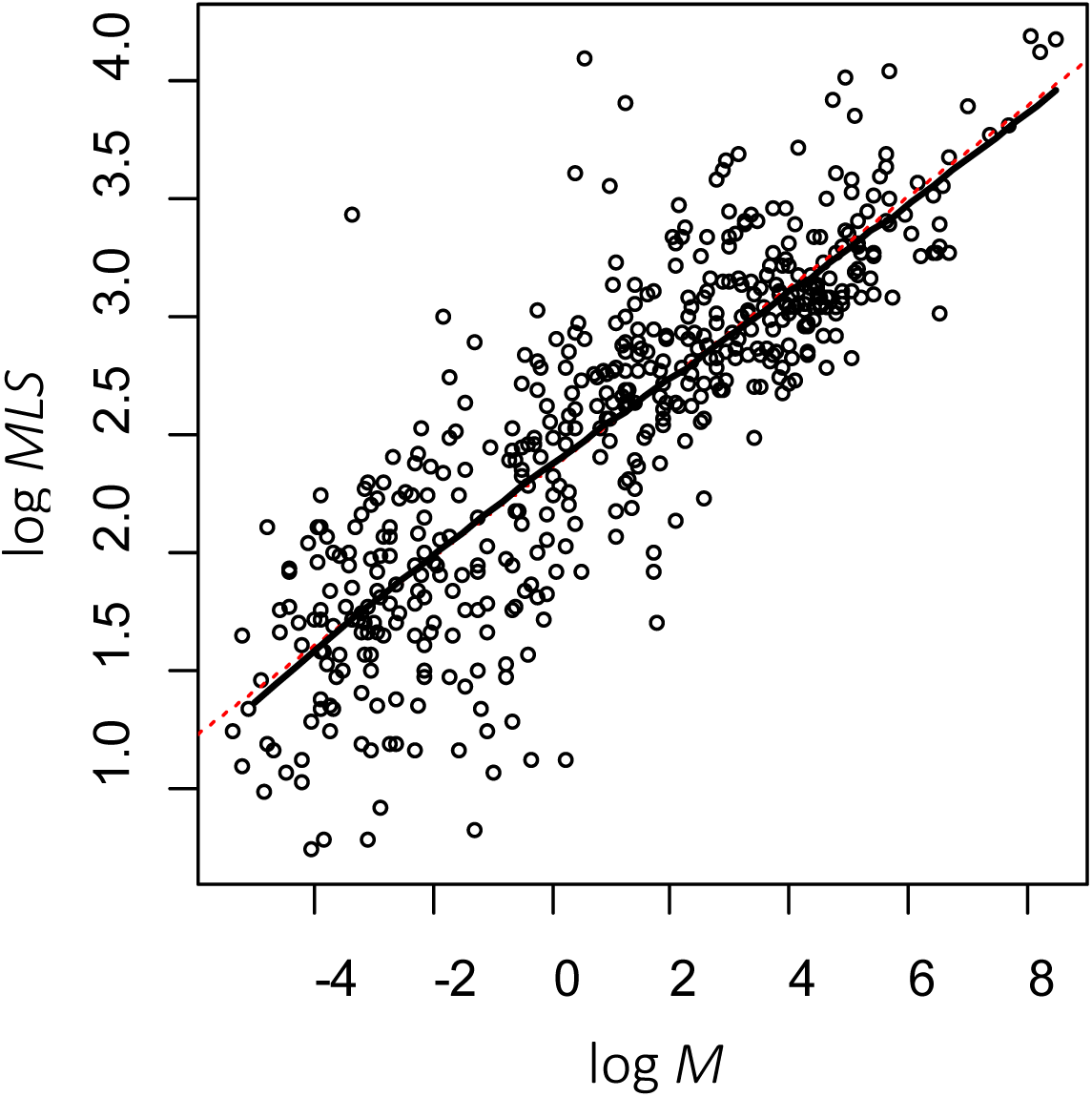
Depicts the fit of terrestrial mammals mass-longevity relationship where *MLS* defined by (6) was calculated using *l*_*x*_ defined by eq. (7) (solid black line). The resulting model is nearly linear as evident by the linear fit (dashed red line).

### Survivorship patterns based on mass

Figure 3A illustrates survivorship curves from eq. (7) for different body sizes generated using *P*_*opt*_ to define the model coefficients by eq. (8). The survivorship curves switch from Type I in large-mass animals to Type II in small-mass animals. The model does not incorporate juvenile mortality so it does not generate Type III survivorship curves characterized by high early-life mortality and lower adult mortality. The shaded area around each curve depicts the variations generated by the SD of parameters (Table 1). Figure 3B illustrates the effect of parameter variability on the *MLS* estimates, which corresponds to curves intersections with the dash line representing *l*_*x*_ = 0.03. The spread of the curves across the line increase approximately linearly with the *MLS* estimate and therefore the error in *MLS* estimate increases in a linear manner with error in *P*_*opt*_. To illustrate, for a 1000 kg mammal the SD variations in *P*_*opt*_ results in a ∼ 1 yr SD on a 50 yr estimate of *MLS* and for a 1 kg mammal the SD is ∼ 0.1 yr on a 10 yr *MLS*. Thus, the ratio of the variability to the mean MLS is ∼ 0.02. In contrast, the SD of the ratio of the mass specific observed *MLS* to the mass-specific mean *MLS* is 0.47. Thus, the variability in the data is a factor of 100 greater than the uncertainty on the predicted *MLS*.

**Fig. 3.**
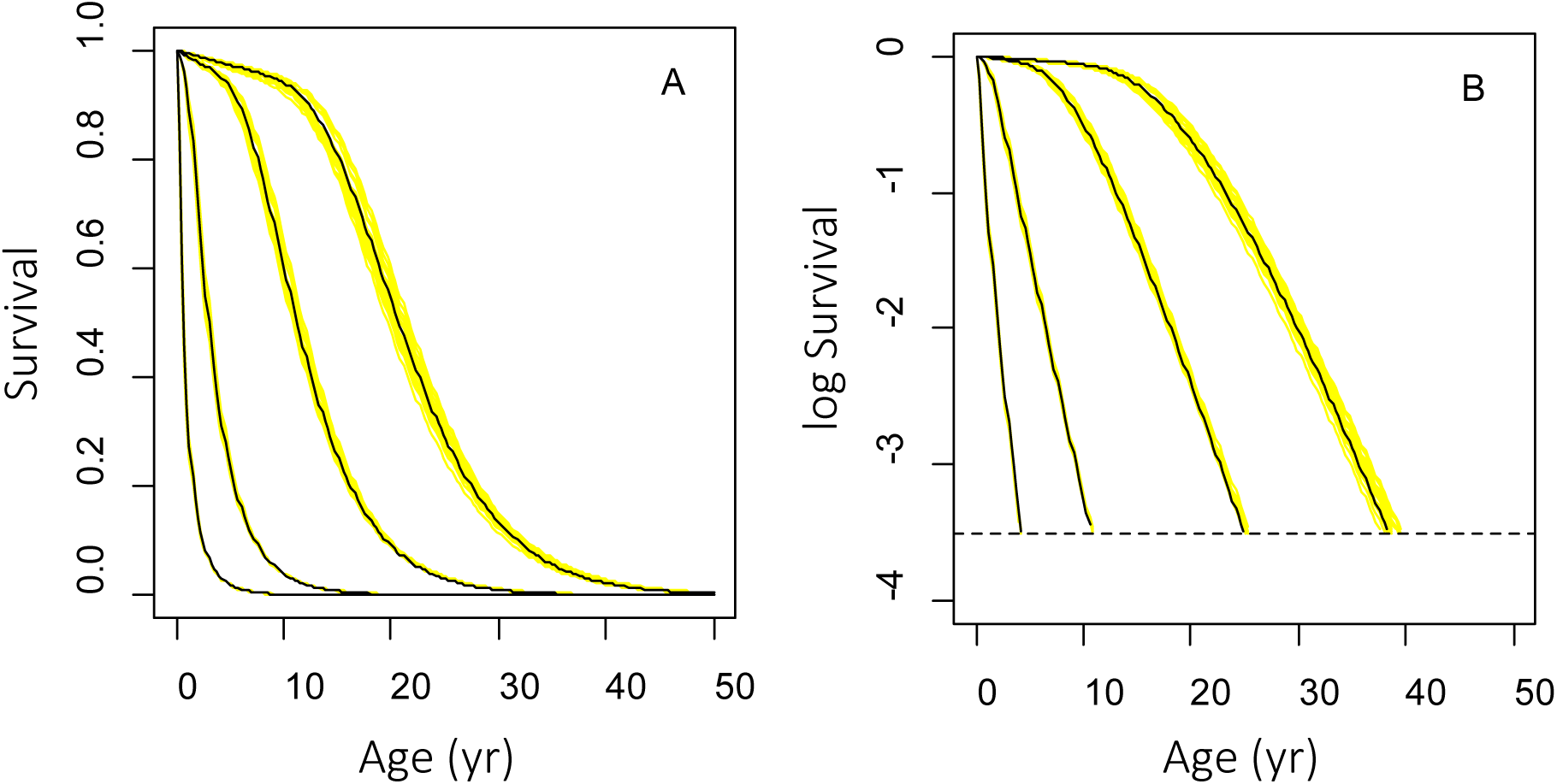
Examples of survivorship curves (A) and log survival (B) for animal masses 0.01, 1 100 and 1000 kg. In B the dashed line depicts the *MLS* of *l* = 0.03. The black lines are model predictions from eq. (7) using the fitted parameters *P*_*opt*_ given in Table 1. The yellow shadow lines illustrate survivorship curves with parameters deviations given in Table 1.

The model partitions mortality into extrinsic and intrinsic processes defined by eq. (5) and eq. (3) respectively. Because in this analysis challenge magnitudes are assumed *β* ≫ 1, the extrinsic survival rate becomes exponential as 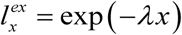. As evident in Fig. 4, survival is dominated by intrinsic processes and effect of extrinsic mortality is mostly evident in early life.

**Fig. 4.**
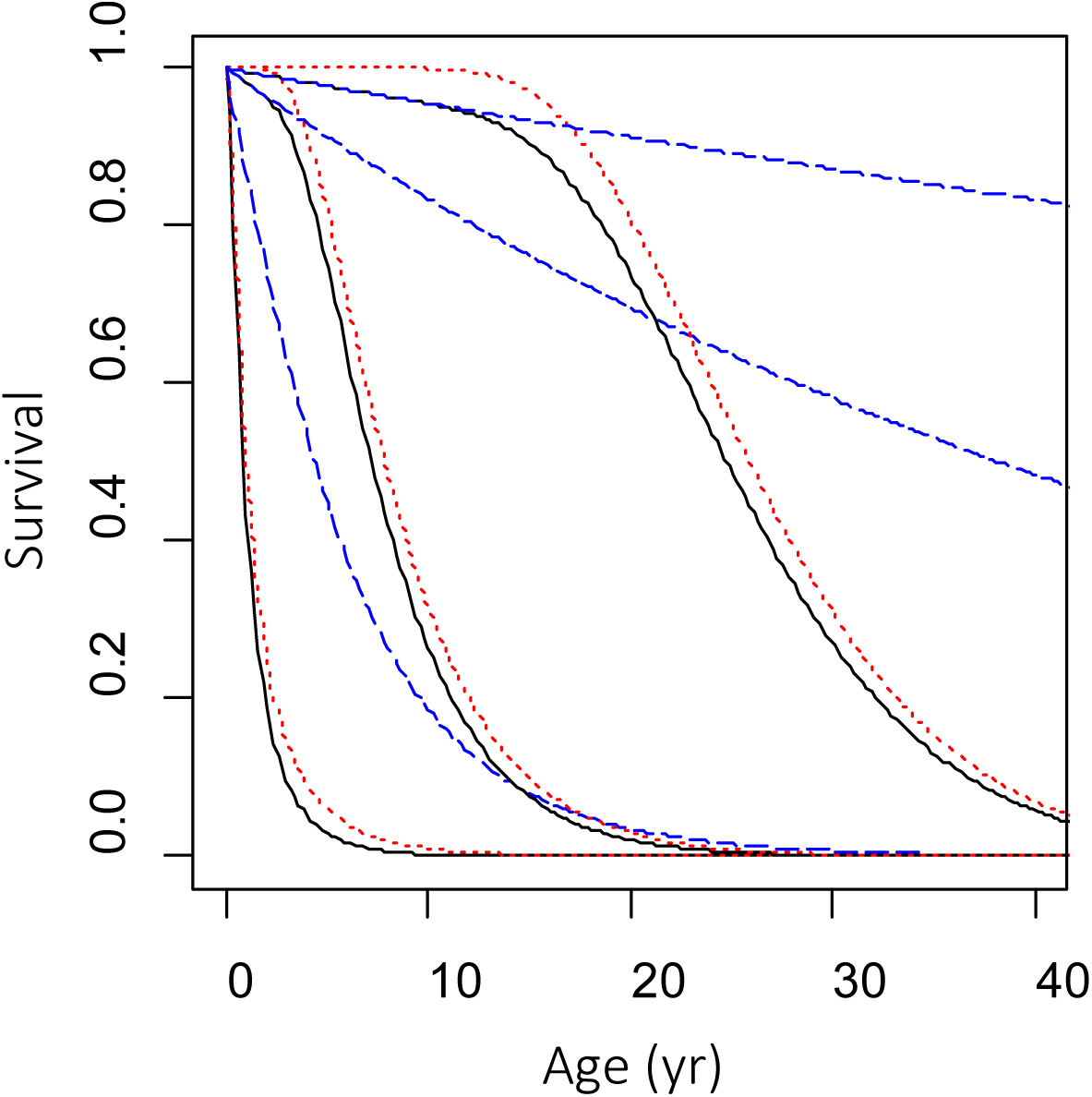
Examples of survivorship (black lines) decomposed into survivorship from extrinsic processes only (dotted red lines) and extrinsic processes only (dashed blue lines). The left most set of lines represents a 0.02 kg animal, the middle grouping a 20 kg animal and the right most grouping a 2000 kg animal.

Figure 5 illustrates the relationship of maximum lifespan to the expected lifespan as the ratio, *MLS/ELS*. The ratio declines with increasing adult body size, reflecting a transition from a Type II survivorship curve to a Type I curve. In effect, the ratio quantifies the increasing rectangularization of the survivorship curve with increasing mass. Importantly, with rectangularization, mortality shifts towards old age. This transition emerges as a property of the mass dependencies of the deterministic and stochastic rates of loss of vitality in eq. (2). To illustrate, note that variance in vitality accumulates with age *x* as Var[*v*_*x*_] ∼ *σ* ^2^ *x* and to a first order the *ELS* is *x* ∼ 1/*ρ*. Then variance in vitality at the *ELS* is Var[*v*_*ELS*_] ∼ *σ* ^2^/*ρ*. Next, using the mass relationships of eq. (8) it follows Var[*v*_*ELS*_] ∼ M^−2*k*^/*M*^−*k*^ = *M*^− *k*^ where *k* ∼ ¼. Thus, small species experience greater variance in vitality over their lifespan than do large species, which is manifested in the shift from Type II to Type I survivorship as body mass increases. Importantly, variations in the extrinsic mortality rate, illustrated in Fig 5 as a change in three orders of magnitude, have small effects on the ratio of maximum to expected lifespans. This suggests that, to a first order, survivorship patterns can be inferred from body size with the environmental effects being secondary.

**Fig. 5.**
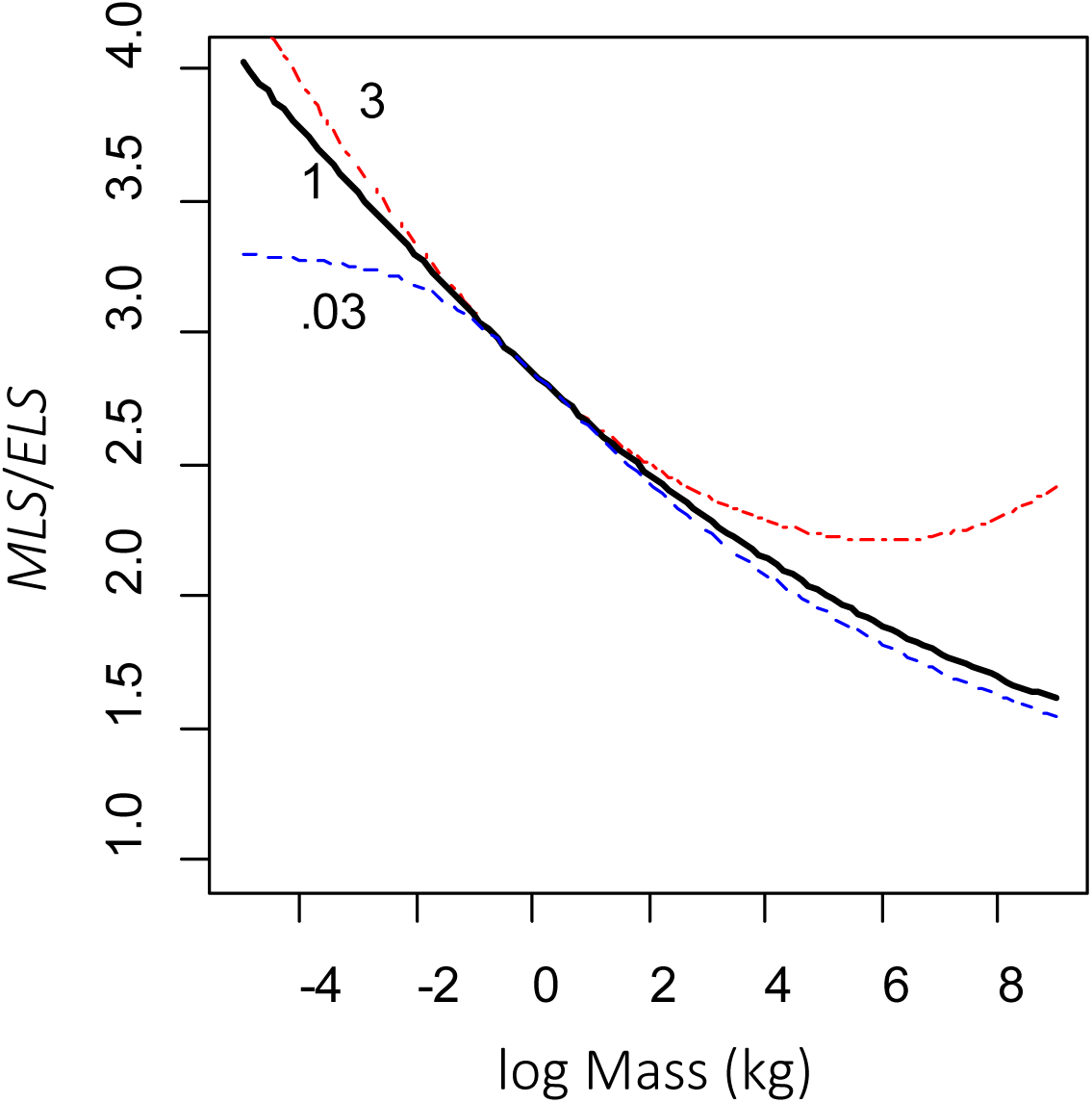
Ratio of maximum lifespan to expected lifespan as a function of the log mass of animal for the *P*_*opt*_ fitting parameters. The red and blue dashed lies on either side of the black line represent changes in the frequency of challenges as 0.03*λ* and 3*λ*.

### Survivorship patterns based on fitting survival data

Figure 6 illustrates the fit of eq. (7) to observed survivorship data representing captive and wild species, different life history strategies and body sizes. For each example, survivorship curves were generated from the fitted *Q*_*S*_ parameters. Figure 6 also depicts survivorship curves derived from *Q*_*M*_, which were generated from eq. (8) using species mass and *P*_*opt*_. Table 2 gives the *Q*_*S*_ and *Q*_*M*_ for each example.

**Fig. 6.**
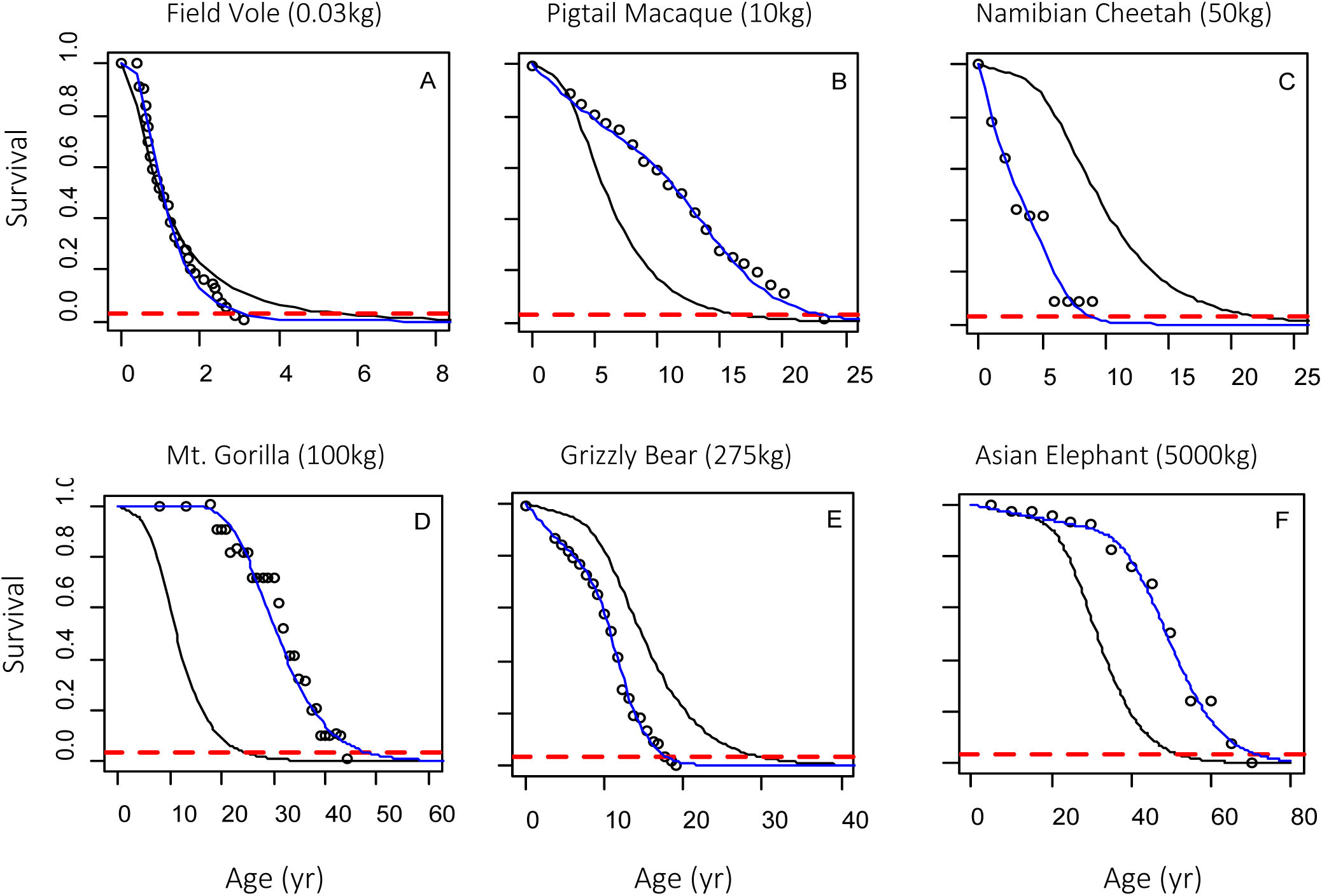
Survivorship data (∘) for example species. The black lines were generated with *Q*_*M*_ parameter sets. The blue lines passing through the observations were generated with the *Q*_*S*_ parameter sets (Table 2). The dashed red line depicts survival of 0.03 so the intersection of a survivorship curve with the line depicts the *MLS*.

**Table 2.** Parameter sets *Q.* = [*ρ.*,*σ.*, *λ.*] to generate survivorship curves in Fig. 6 using eq. (7). Sets *Q*_*S*_ are estimated by fitting eq. (7) to published survival data. Sets *Q*_*M*_ are derived from eq. (8) using animal mass and fitting parameters *P*_*opt*_. Early life mortality was removed from the survival data by linearly projecting survival to zero age using the survivorship curve slope at onset of adult age.

| Example species | Mass (kg) | Adult age (yr) | Parameters and factors | $\rho$ (yr<sup>-1</sup>) | $\sigma$ (yr<sup>-1</sup>) | $\lambda$ (yr<sup>-1</sup>) |
| --- | --- | --- | --- | --- | --- | --- |
| Field Vole | 0.03 | 0.17 | $Q_M$ | 0.54 | 0.83 | 0.127 |
| | | | $Q_S$ | 0.81 | 0.52 | 0 |
| Pigtail Macaque | 10 | 2 | $Q_M$ | 0.132 | 0.206 | 0.022 |
| | | | $Q_S$ | 0.060 | 0.065 | 0.048 |
| Namibian Cheetah | 50 | 0 | $Q_M$ | 0.09 | 0.14 | 0.014 |
| | | | $Q_S$ | 0.14 | 0.10 | 0.22 |
| Mountain Gorilla | 100 | 8 | $Q_M$ | 0.076 | 0.118 | 0.011 |
| | | | $Q_S$ | 0.032 | 0.044 | 0.0 |
| Grizzly Bear | 275 | 5 | $Q_M$ | 0.06 | 0.093 | 0.008 |
| | | | $Q_S$ | 0.078 | 0.065 | 0.036 |
| Asian Elephant | 4000 | 5 | $Q_M$ | 0.03 | 0.046 | 0.003 |
| | | | $Q_S$ | 0.019 | 0.027 | 0.003 |

### Comparison of mass-based and observation-based survivorship curves

Comparison of survivorship curves generated with the *Q*_*S*_ and *Q*_*M*_ parameter sets illustrate the degree that survival can be predicted from body size. For this discussion, consider the *Q*_*S*_ parameters represent the actual processes shaping the observed survival and the *Q*_*M*_ parameters as first order estimates based on an aggregate of species masses. The ratios *Q*_*S*_/*Q*_*M*_ (Table 3) provide information on the limitations of the mass-based estimates of survival as well as reasons for their biases. In general, the *Q*_*M*_ parameters underestimate survival in the macaque, gorilla and elephant, over predict survival for cheetah and bear and fit the vole data relatively well.

**Table 3.** Ratios of model parameter sets *Q*_*S*_ to *Q*_*M*_.

| Example species | $\rho_S/\rho_M$ | $\sigma_S/\sigma_M$ | $\lambda_S/\lambda_M$ |
| --- | --- | --- | --- |
| Field Vole | 1.5 | 0.7 | 0.0 |
| Pigtail Macaque | 0.5 | 0.3 | 2.2 |
| Namibian Cheetah | 1.5 | 0.7 | 16.0 |
| Mountain Gorilla | 0.4 | 0.4 | 0.0 |
| Grizzly Bear | 1.3 | 0.7 | 4.4 |
| Asian Elephant | 0.7 | 0.6 | 0.9 |

#### Field Vole

For the field vole example (Fig. 6A), *Q*_*M*_ fits the survival data relatively well over most ages but over predicts *MLS*, as illustrated by the difference in ages at which the *Q*_*M*_ and *Q*_*S*_ curves crosses the dashed representing *l* = 0.03; a difference of 3 years. Notably the *Q*_*S*_ set has *λ* = 0 (Table 3). This prediction of no environmental challenges comports with the voles being individually weened and held in single cages with occasional handling and testing (Selman et al. 2008). The captive conditions also may in part account for the increased rate of vitality loss due to stress. However, the variability in rate of vitality loss for *Q*_*S*_ was lower than estimated with *Q*_*M*_ (Table 3), perhaps resulting from the uniformity of stress.

#### Pigtail Macaque

For the pigtail macaque example (Fig. 6B), the survival data was derived from the males in the Washington Regional Primate Research Center Pigtail Macaque Colony (Ha et al. 2000). The intrinsic parameters for *Q*_*S*_ were both lower than the *Q*_*M*_ parameters, indicating the rate of loss of vitality for the primate was about half that projected from *Q*_*M*_ (Table 3). Notably, the infant mortality in the population was high and therefore the surviving population may reflect a selection of stronger individuals. Additionally, the population was under sophisticated veterinary and husbandry care. These factors together could account for lower vitality rate parameters. The rate of extrinsic mortality was higher by a factor of two. Physical injury under captivity possibly was greater than would be predicted by the aggregate from the AnAge data.

#### Namibian Cheetah

The difference in the cheetah survivorship curves derived from *Q*_*S*_ and *Q*_*M*_ (Fig 6C) resulted from significant differences in extrinsic mortality (Table 3). The higher *λ*_*S*_ can be attributed to the 90% overlap of the cheetah habitat with commercial farms in Namibia. As a consequence of this overlap, over the period of the study (1991-2000) between 100 and 300 animals were removed by farmers yearly (Marker et al. 2003).

#### Mountain Gorilla

The survivorship data for female mountain gorilla (Fig. 6D) represents a small population in the Virunga Volcano region of Rwanda, Uganda, and Democratic Republic of Congo. The *Q*_*S*_ parameters compared to *Q*_*M*_ parameters had significantly lower mean and variance in the rate of loss of vitality and no extrinsic mortality (Table 3). The biological reasons for these differences are unclear but a number of caveats are worth noting. The population had experienced significant poaching in the past but during the study the level was low (Robbins et al. 2011). The high longevity of the gorilla population relative to the mass-based prediction reflects a similar pattern for the pigtail macaque population. One possible explanation involves lower stress levels in primates compared to other species in the AnAge database. For example, studies with baboons indicated that the level of early adversity could affect lifespan by a factor of two. Under high stress, baboon *MLS* was < 15 yrs, while under low stress conditions, *MLS* was >25 yrs (Tung et al. 2016). Notably, for both the macaque and gorilla, *MLS* based on *Q*_*S*_ and *Q*_*M*_ were different by about 15 yrs. The lower estimated rate of vitality loss in the primate populations may also involve metabolic differences relative to the majority of species in the AnAge database.

#### Grizzly Bear

The survivorship curve (Fig. 6E) for grizzly bear was extracted from Figure 7 in (Knight and Eberhardt 1985) depicting the grizzly bear population in Yellowstone National Park in the 1970ies. During this time, the population was under significant hunting pressures at the park boundaries and illegal hunting within the park. Compared to *Q*_*M*_ the *Q*_*S*_ parameters had higher rates of extrinsic mortality and vitality loss while the vitality rate variance was lower (Table 3). Thus, the mass based predictions did not capture the actual population’s higher rate of aging (*ρS* > *ρM*) or higher rate of extrinsic mortality (*λS* > *λM*).

#### Asian Elephant

The survivorship for Asian elephants (Fig 6.F) was derived from males in a captive breeding population in the Mudumalai and Anamalai Wildlife Sanctuaries of India (Sukumar et al. 1997). The *Q*_*S*_ parameters were uniformly lower than the *Q*_*M*_ parameters resulting in the mass-based prediction underestimating *MLS* by a decade. This difference may in part reflect the protected habitat of the elephants relative to the aggregate of the AnAge species. Additionally, the model may overestimate the rate of aging in large species.

## Discussion

This paper presents a model linking metabolic and vitality theories to describe the effect of body size on intrinsic and extrinsic rates of mortality. The result is a quantitative framework for exploring the contributions of body size on physiological and environmental factors that shape survivorship. Because physiological time scales of animals have allometric relationships with body size, the framework should provide more resolution of the factors that shape animal fitness and evolution. In particular, a future extension of this model will explore environmental and physiological processes underlying Cope’s rule.

The model is developed from a dimensional premise linking the replicative capacity of HSC and body-mass regulated metabolism (Gillooly et al. 2012). Hematopoiesis is focal because through asymmetrical differentiation, HSCs renew and produce progenitor cells that follow different pathways to daily generate the 10^11^-10^12^ new blood cells required for oxygen transport and immune functions of an organism. The hypothesis that HSC replicative senescence limits animal longevity rests on the observation that mammals have about the same number of HSC. Thus, to a first order, the maximum lifespan of an animal is determined by the ratio of the number of HSC divided by their rate of replication. Because cell replication rate is proportional to the ¼-power of body size, longevity should have a similar relationship to body size. While there is no direct evidence that the HSC pool is under simple replicative control nor limits lifespan, the underlying premise that cell divisions result in cumulative damage that limits system function is supported by studies of hematopoiesis aging (Park 2017). Thus, the replicative limit hypothesis provides a useful framework on which to develop a cell-based model linking body size to survivorship while avoiding details of aging.

The properties of replicative senescence are readily described by vitality theory in which the intrinsic mortality rate is determined by the stochastic arrival time of vitality (a measure of the HSC pool) to a zero boundary (the Hayflick limit) (Anderson et al. 2017; Li and Anderson 2009; Li and Anderson 2013). Relating the rate of vitality decline to body size with metabolic theory (West et al. 1997) yields a model for the pattern of survivorship from intrinsic processes only. Additionally, a macro ecological study between predator-prey interactions and body size (Hatton et al. 2015) provides a measure of the extrinsic mortality rate. Combining the two rates, a base effect of body size on intrinsic and extrinsic survivorship patterns is characterized by five parameters: two allometrically rescale of body size effects and three proportionally scale the rescaled body size to the intrinsic and extrinsic rate parameters. The parameters are estimated by fitting the model to data on maximum life span vs. adult body size of animals ranging from grams to tons.

Several properties emerge from the model. Firstly, the partition between intrinsic and extrinsic mortality changes with body size, with extrinsic mortality becoming more important at smaller sizes. This shift provides a biological mechanism for survivorship curves in which mortality occurs mostly in late life (Type I curve) or is constant over life (Type II curve). Intrinsic mortality, which occurs in old age, is characteristic of large body animals that have Type I survival. Intrinsic mortality, which has the greatest effect in early life, is characteristic of small animals commonly exhibiting Type II survival. These patterns emerge solely because of the stochastic properties of vitality loss, or by inference, the stochastic nature of hematopoiesis aging.

In fitting the model to *MLS* data across a wide range of body masses the resulting parameters represent a generalized partition of intrinsic and extrinsic processes shaping survivorship. To explore cohort-level differences in the partition, the vitality model was fit to survivorship profiles from six species from field voles (0.01 kg) to elephants (5000 kg). A comparison of the parameters derived using only mass *Q*_*M*_ to parameters derived by fitting the survivorship data *Q*_*S*_ provides insights to the possibilities and limitations of inferring survivorship from mass information only.

Firstly, the model fit the survivorship data very well and so the *Q*_*S*_ parameters are, to the degree possible with the model, accurate representations of the intrinsic and extrinsic processes shaping the observed survival. The ratios of the *Q*_*S*_ to *Q*_*M*_ parameters identifies habitat-specific and life history factors that are not captured in the mass-derived survivorship curves. The clearest patterns involve habitat-specific effects on the extrinsic mortality rate. For species in controlled laboratory or protected habitats, the *Q*_*S*_ extrinsic mortality rates were lower than the *Q*_*M*_ estimates. In contrast, the opposite pattern was observed for species in wild habitats, with the difference reflecting the amount of hunting pressure on the species.

A second result involves the rate of vitality loss, which is assumed to reflect the rate of aging in hematopoiesis. For the two primate examples, the vitality loss rate was significantly lower in the *Q*_*S*_ estimates. This presents the intriguing possibility that the rates of aging in primates are lower than occurs in nonprimates of similar size. The ability of the model to separate intrinsic mortality from total mortality highlights the unique properties of primates; a result most evident for humans.

A third result is the agreement of the fitted allometric coefficients with theory. Fitting log *MLS* against log *M* over 6 orders of magnitude yielded the metabolic coefficient of *h* = 0.24 with a variation of 0.003 and the macroecology coefficient of *k* = 0.30 with a variation of 0.013. Alternative forms of metabolic theory suggest the metabolic coefficient lies between 1/4 ≤ *k* ≤ 1/3 (e.g. Darveau et al. 2002; Dodds et al. 2001; Kleiber 1947; West et al. 1997) while an aggregate for the macroecology coefficient is *k* = 0.3 (Hatton et al. 2015). Thus, the model estimates of the coefficients from lifespan data are essentially equivalent to the range suggested in the literature. Furthermore, the difference 0.24 vs. 0.25 or 0.33 for *h* does not affect the properties of the model. Thus, the model dimensions can be reduced to three parameters by setting the allometric parameters to their theoretical values *h* = 0.25 and *k* = 0.3.

Combining metabolic and vitality theories extends both in new directions. It points to a more mechanistic foundation for vitality and extends the metabolic theory into stochastic mechanisms of population survival. A merged theory illustrates a quantitative framework to explore the interactions of physiology and environment in shaping population fitness as well as the effects of aging across the biomass spectrum of animals.

## Acknowledgements.

This work was supported by National Institute of Health Grant R21AG046760.

## Footnotes

1 A healthy body of literature has discusses where the coefficient is between 2/3 and ¾ (Banavar et al. 2010; da Silva et al. 2006; Dodds et al. 2001; Lee 2015) For the model here the value of the exponent is not germane.

2 Leukocytes, the central actors of the immune system, are produced from hematopoietic stem cells in the bone marrow and the lymph system. The leukocytes circulate as a component of blood to detect and remove pathogens from the body and repair damaged tissue. The gradual reduction of immune system function with age involves essentially all the aging hallmarks but of particular importance are the loss of proteostasis and the reduction in the number of naïve T cells and stem cells with age (Ponnappan and Ponnappan 2011). Collectively these processes are known as immunosenescence (Weiskopf et al. 2009) and result in inflammaging, both of which are hallmarks of aging leading to death (Franceschi et al. 2007). While immune system function is an index of aging, degradation of other processes working across the molecular, cellular and system levels are also important (López-Otín et al. 2013).

3 This is an upper limit of cell replication, determined by the attrition of the protective telomeres end caps on chromosomes (Hayflick and Moorhead 1961). Once the telomere length reaches it critical length cell replication stops. However, telomerase can be maintain TL (Blackburn et al. 2015). Evidence suggests that TL declines with lifespan in some species but not others. Telomerase varies with body size but not life span (Monaghan 2010).

